# Phylogenetically novel uncultured microbial cells dominate Earth microbiomes

**DOI:** 10.1101/303602

**Authors:** Karen G. Lloyd, Joshua Ladau, Andrew D. Steen, Junqi Yin, Lonnie Crosby

**Affiliations:** Department of Microbiology, University of Tennessee, Knoxville, TN 37996; Gladstone Institutes, University of California San Francisco, San Francisco, CA 94158; Department of Earth and Planetary Sciences, University of Tennessee, Knoxville TN 37996; Joint Institute for Computational Sciences, University of Tennessee, Knoxville, TN 37996

## Abstract

To unequivocally determine a microbe’s physiology, including its metabolism, environmental roles, and growth characteristics, it must be grown in a laboratory culture. Unfortunately, many phylogenetically-novel groups have never been cultured, so their physiologies have only been inferred from genomics and environmental characteristics. Although the diversity, or number of different taxonomic groups, of uncultured clades has been well-studied, their global abundances, or number of cells in any given environment, have not been assessed. We quantified the degree of similarity of 16S rRNA gene sequences from diverse environments in publicly-available metagenome and metatranscriptome databases, which we show are largely free of the culture-bias present in primer-amplified 16S rRNA gene surveys, to their nearest cultured relatives. Whether normalized to scaffold read depths or not, the highest abundance of metagenomic 16S rRNA gene sequences belong to phylogenetically novel uncultured groups in seawater, freshwater, terrestrial subsurface, soil, hypersaline environments, marine sediment, hot springs, hydrothermal vents, non-human hosts, snow and bioreactors (22-87% uncultured genera to classes and 0-64% uncultured phyla). The exceptions were human and human-associated environments which were dominated by cultured genera (45-97%). We estimate that uncultured genera and phyla could comprise 7.3 × 10^29^ (81%) and 2.2 × 10^29^ (25%) microbial cells, respectively. Uncultured phyla were over-represented in meta transcript omes relative to metagenomes (46-84% of sequences in a given environment), suggesting that they are viable, and possibly more active than cultured clades. Therefore, uncultured microbes, often from deeply phylogenetically divergent groups, dominate non-human environments on Earth, and their undiscovered physiologies may matter for Earth systems.

## Introduction

Direct sequencing of environmental DNA has shown that most microbial lineages have not been isolated in pure culture (1–3). However, cellular abundances and viability states of uncultured microbes at different levels of phylogenetic divergence from their closest cultured relative are unknown. Cellular abundance and viability may, in some cases, signify importance to current ecosystem functions, as opposed to members of the rare biosphere which become important for ecosystem functioning when conditions change (4). With the exception of keystone species which can have great ecosystem importance even at low biomass concentrations, prokaryotic abundance and viability is generally an indicator for participation in current ecosystem functions (1).

Quantifying the cellular abundance of all microbial taxa in any sample is challenging. Fluorescent in situ hybridization (FISH) allows fluorescent tagging of a taxonomic group whose cells can then be counted under a microscope (2). However, FISH requires developing probes for phylogenetic groups one-by-one, which is impractical for quantifying highly diverse natural samples that are often comprised of thousands of species (3). Furthermore, FISH techniques are not always quantitative in all environments, due to taxon-specific biases in probe efficacy (4, 5). Quantitative PCR has the same low-throughput limitations, since individual measurements must be made for each taxon, and primer bias makes them not absolutely quantitative (4). However, understanding the total cellular abundance of uncultured clades of archaea and bacteria in all environments on Earth is an essential question in microbiology, so we approximated it using the data available in public databases.

Genes encoding the 16S rRNA small subunit of the ribosome are the most commonly-used taxonomic and phylogenetic identifier for bacteria and archaea, and most scientific journals make publication contingent on the posting of 16S rRNA gene sequences to public databases. Therefore, the National Center for Biotechnology Information (www.ncbi.nlm.nih.gov) houses a nearly-complete database of full length 16S rRNA gene sequences. This database is subject to biases since the gene entries have undergone exponential amplification from their initial abundances, and small mismatches between DNA primers and different taxa are magnified during this amplification (5). However, we examined it here since it incorporates microbial phylogenetic information from thousands of different research studies. Assembled metagenomes provide a less-biased accounting of 16S rRNA genes from a given environment. Here all DNA is chemically extracted from a sample, purified, sequenced in a small-read high throughput platform, and then bioinformatically assembled to contigs. Full length 16S rRNA genes can be identified in these contigs using hidden Markov model-based programs like RNAmmer (6). If the sequencing depth is great enough, quantifying read recruitment to each 16S rRNA gene provides the closest approximate quantification of individual 16S rRNA genes currently available.

Cellular activity, however, is as important to environmental functions as cellular abundance (1). In cultured cells, rRNA content correlates with cellular activity (18), although no universally predictive relationship between those two parameters has been identified (19). Metatranscriptomes, in which 16S rRNA transcripts are converted to cDNA and sequenced without the use of primers, provide an estimate of which cells contained ribosomes, and therefore were at least poised for activity in the environment (19).

We determined the percent similarity of nearly all 16S rRNA gene sequences from public databases, to get a first estimate of the global abundance of microbial clades at different levels of similarity to their nearest cultured relative in different environments. The metagenomic and metatranscriptomic datasets show that uncultured clades dominate cellular abundance of non-human Earth environments. Knowing the global abundance of cells from uncultured taxa is crucial for estimating the importance of uncultured lineages to ecosystem functions, determining the appropriateness of using cultured microbes as model systems for natural environments, and predicting the causes of unculturability.

## Materials and Methods

Primer-amplified sequences were obtained from www.arb-silva.de Silva123Ref (5), which contains chimera-checked, high quality, >900 bp (for archaea) and >1200 bp (for bacteria) 16S rRNA gene sequences, almost all of which were Sanger-sequenced clone inserts from primer-amplified PCR products. This yielded 952,509 bacterial and 51,608 archaeal sequences from 4,743 studies that employed a wide variety of primers. Genes annotated as 16S rRNA and >900bp were collected from the Joint Genome Institute IMG/M for metagenomes larger than 1 GB total or metatranscriptomes larger than 60 Mb total (6). Too few metatranscriptomes were available from humans, human-adjacent environments, rock, snow, hydrothermal vents, hypersaline environments, or marine sediments to be included. Scaffold read depths were available for metagenomes, but not metatranscriptomes.

Metagenomes and metatranscriptomes are prone to chimera production during assemblies of short reads along the highly conserved 16S rRNA gene (7). We therefore implemented uChime (8) in mothur (9) with the Silva Gold alignment to identify and remove a further 1.3% and 0.6% of possible chimeras from metagenomes and metatrancriptomes, respectively. Further chimera checks are described below. Taxonomic identifications were made for each sequence in the metagenomic and metatranscriptomic datasets in mothur (9) for alignment, pre-clustering, and classification to silva.nr_v132

(10) Sequences identifying as chloroplasts, mitochondria, or eukaryotes (<1% of sequences) were removed.

BLASTn was used to determine the percent identity of each sequence to its single most-closely related 16S rRNA gene sequence from cultured archaea (4,170 sequences) or bacteria (22,150 sequences) obtained from Arb-Silva. Only cultured archaea and bacteria with official names from the International Journal of Systematic Bacteriology or the International Journal of Systematic and Evolutionary Microbiology were included, exc_ludi_n_g_ candidatus organisms or enrichments. Rather than relying on annotations to separate archaea and bacteria in metagenomes and metatrancriptomes, sequences were queried against a database with bacteria and archaea combined to get the top hit. We used a BLASTn implementation parallelized for high performance computation, HPC-BLAST(11), on the Beacon cluster (12) at the Joint Institute for Computational Sciences. The alignment results of HPC-BLAST are compatible with those of NCBI BLAST.

A few metagenomic and metatranscriptomic 16S sequences did not yield BLASTn hits, so were not considered further. For sequences with query alignment lengths <300 bp, percent identity increased with decreasing alignment length, suggesting that these were partial hits to small conserved regions, so they were removed from the analysis. Short query alignment lengths could also signify chimeras. Therefore sequences with < 90% alignment length to their closest cultured relative were aligned with BLASTn to the SilvaNR database, containing environmental DNA sequences. Sequences with <90% alignment to both the cultured and Silva NR databases were considered to be chimeric and were removed from analysis. This removed 6% of the metagenomic database, leaving 39,426 bacterial and 13,404 archaeal sequences from 1,504 metagenomes, as well as 7% of the metatranscriptomic database, leaving 9,396 bacterial and 3,863 archaeal sequences from 381 metatranscriptomes. Each remaining sequence was manually categorized into one of 14 environment types, based on user-provided metadata (Tables S1 and S2).

16S rRNA gene sequences that shared more than 96.6% sequence identity with a cultured organism were considered to be in the same genus, and sequences that shared at least 86% sequence similarity were considered to be in the same phylum (13). These create “similarity bins” of cultured species to genus, uncultured genus to class, and uncultured phyla and higher. For primer-amplified, metagenomic, and metatranscriptomic datasets, the fraction of sequences in each similarity bin was calculated for a given environment. In metagenomes for which sequence read depth was available, the fraction in each similarity bin was calculated as the sum of sequence read depths for each similarity bin within each metagenome. These values were averaged for all metagenomes in each environment.

## Results and Discussion

More than a third of primer-amplified 16S rRNA gene sequences were from the same species or genus as a culture (37% for bacteria and 34% for archaea, Fig. 1), in agreement with previous findings that primer-amplified databases skew toward cultured organisms (14–16). However, even in the primer-amplified dataset, the majority of sequences were from uncultured genera or higher taxonomic groups, including 17% and 44% from uncultured phyla in bacteria and archaea, respectively. This suggests that, as a group, uncultured microbes, including those that are very highly divergent, are fairly abundant when all full length 16S rRNA genes in public databases are considered. Metagenomes had lower fractions of 16S rRNA gene sequences from cultured species (Fig. 1), with 15% for both bacteria and archaea based on total sequences and 28% for bacteria and 31% for archaea based on scaffold read depths. The rest of the 16S rRNA gene sequences were from uncultured genera and higher taxonomic groups, with about a third of total sequences from uncultured phyla (36% and 46% without read depths, 24 and 33% with read depths for bacteria and archaea). However, we recognize that it is impossible to absolutely link 16S identity to taxonomic level, since phylogenetic difference is inconsistently related to 16S sequence difference across lineages (13). Therefore, these sequence similarity cutoffs are proxies for degrees of phylogenic novelty rather than rigidly-defined taxonomic levels. By using published values for similarity bins (13), our findings are comparable to other studies and serve as an estimate for phylogenetic novelty informed by the available data. Therefore, 16S rRNA gene sequences from uncultured cells were more abundant than those from cultured cells, suggesting that uncultured microbial clades are not collectively relegated to the “rare biosphere” (17), but are instead the most numerically dominant cells in the public databases.

**Figure 1.**
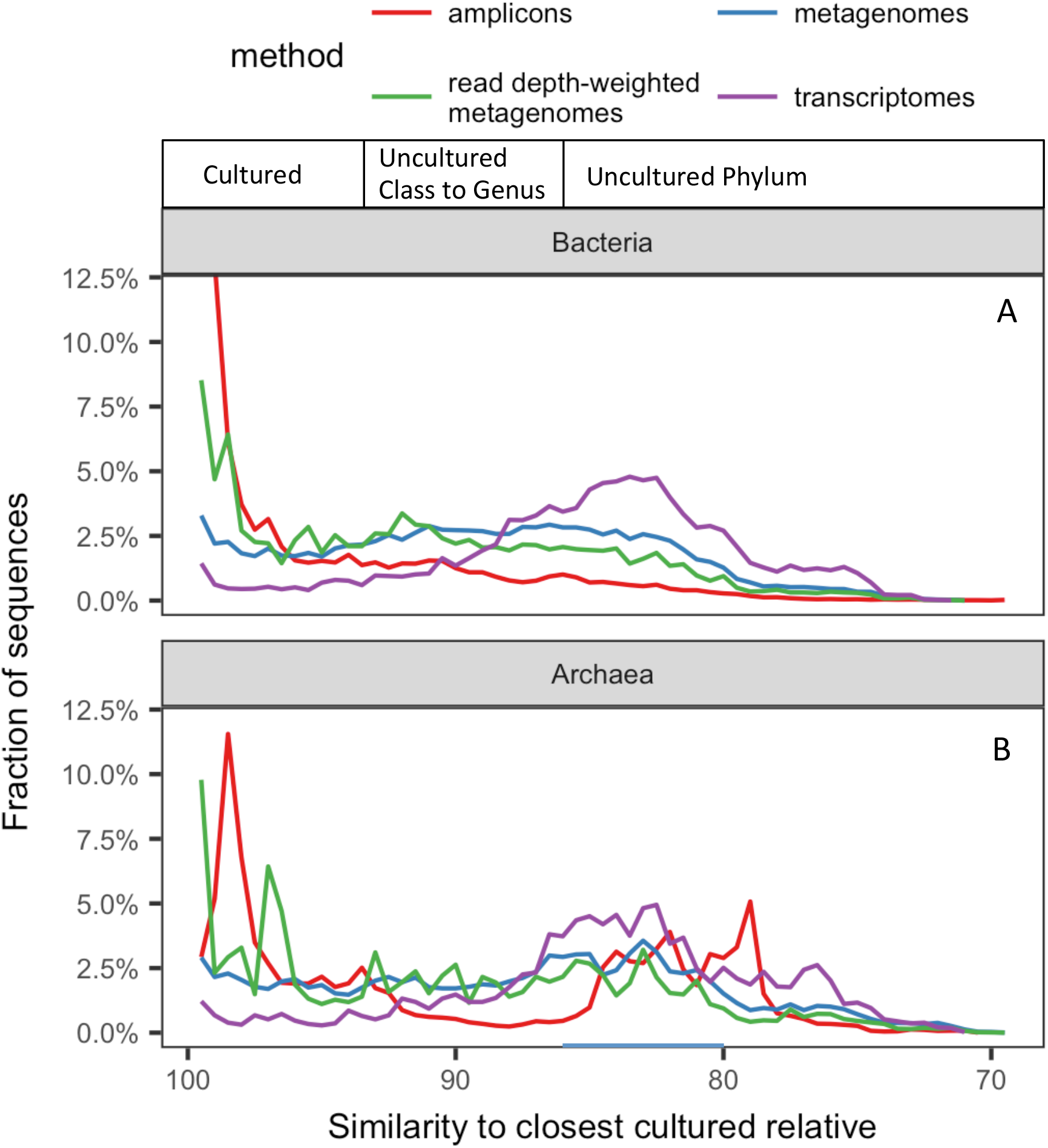
Fraction of 16S rRNA genes from bacteria (A) and archaea (B) in public databases from primer-amplified, metagenomes (with and without read depths), and metatranscriptomes at different percent identities with their closest cultured relative. Box along the top shows estimated cut-offs for different taxonomic level of novelty relative to all cultures (13). Primer-amplified bacterial sequences go up to 30% at 100%) similar to their closest cultured relative, but were removed for clarity.

We found that highly divergent uncultured sequences were better represented in metatranscriptomes than in metagenomes, with only 4% (bacteria) and 5% (archaea) of total sequences from cultured species to genera, and 65% (bacteria) and 71% (archaea) of total sequences from uncultured phyla. Therefore, cells from highly divergent uncultured groups were alive *in situ,* and may even be more active in natural samples than cells from cultured species and genera. A comparison between metagenomes and metatranscriptomes both derived from the same samples in the Gulf of Mexico showed that uncultured clades were indeed active relative to cultured clades (20).

Contributions from uncultured clades varied by environment (Fig. 2). The only environments dominated by sequences from cultured species and genera were the human body and human-adjacent environments (Fig. 2). This result was not due to primer bias, since primer-amplified and metagenomic datasets contained mostly cultured species and genera (45-97%, inclusive of bacteria and archaea). High culturability in human environments likely benefits from a high frequency of culturing efforts, since all culturing happens in the vicinity of humans, and since the study of human diseases has driven much research (21). Uncultured clades were also present in humans and human-adjacent environments, but very few were uncultured at a taxonomic cut-off above family level.

**Figure 2.**
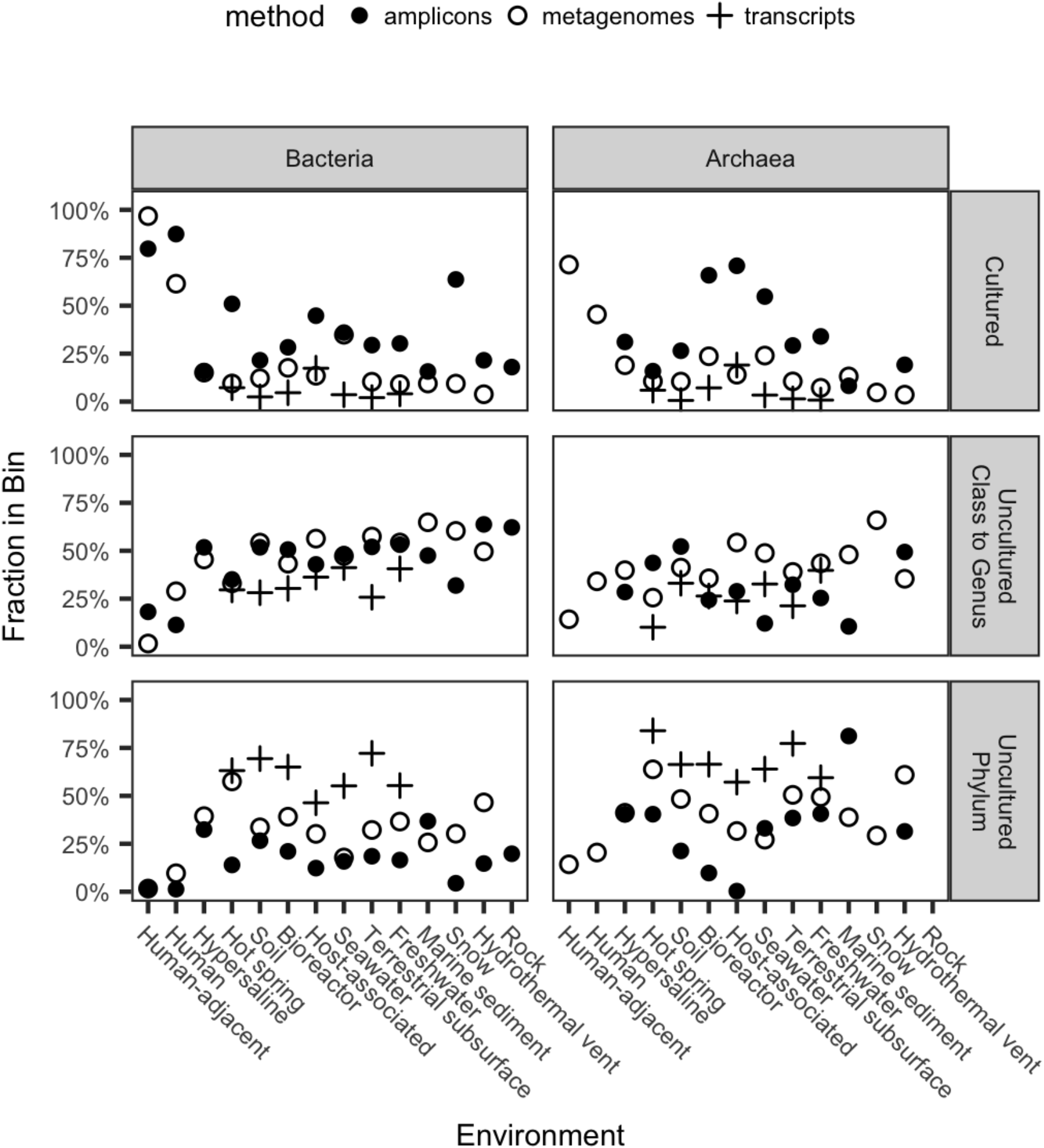
Proportion of 16S rRNA sequences in each category of phylogenetic novelty relative to cultures for each environment, by amplicons, metagenomes (without scaffold read depth), and metatranscriptomes. Closed circles are primer-amplified amplicons, open circles are metagenomes, and crosses are transcriptomes.

Primer bias toward cultures was more severe in all other environments, where uncultured archaea and bacteria were much more abundant in metagenomic datasets than in primer-amplified datasets (Fig. 2). The exception was archaea in marine sediments, possibly indicating that commonly-used primers have good matches to the uncultured phyla that are abundant in these environments (22). To avoid primer bias and account for a high environmental abundance of closely related sequences, we used the metagenomic datasets with read depths to estimate quantifications (Fig. 3). Hypersaline environments were the next best-cultured environments after human environments, with nearly half of archaea and bacteria from cultured genera, and very few from uncultured phyla (Fig. 3). The next best-cultured group was archaea in bioreactors. All other environments had more sequences from uncultured phyla than from cultured genera. Hot springs and hydrothermal vents, in particular, had high frequencies of uncultured phyla in both bacteria and archaea. Even though human host environments were dominated by cultured groups, non-human hosts had as few sequences from cultured archaea and bacteria as did soil, seawater, freshwater, marine sediment, terrestrial subsurface, snow and bioreactors (for bacteria). This suggests that highly divergent uncultured microbes, possibly with novel functions, dominate non-human environments on Earth.

**Fig. 3.**
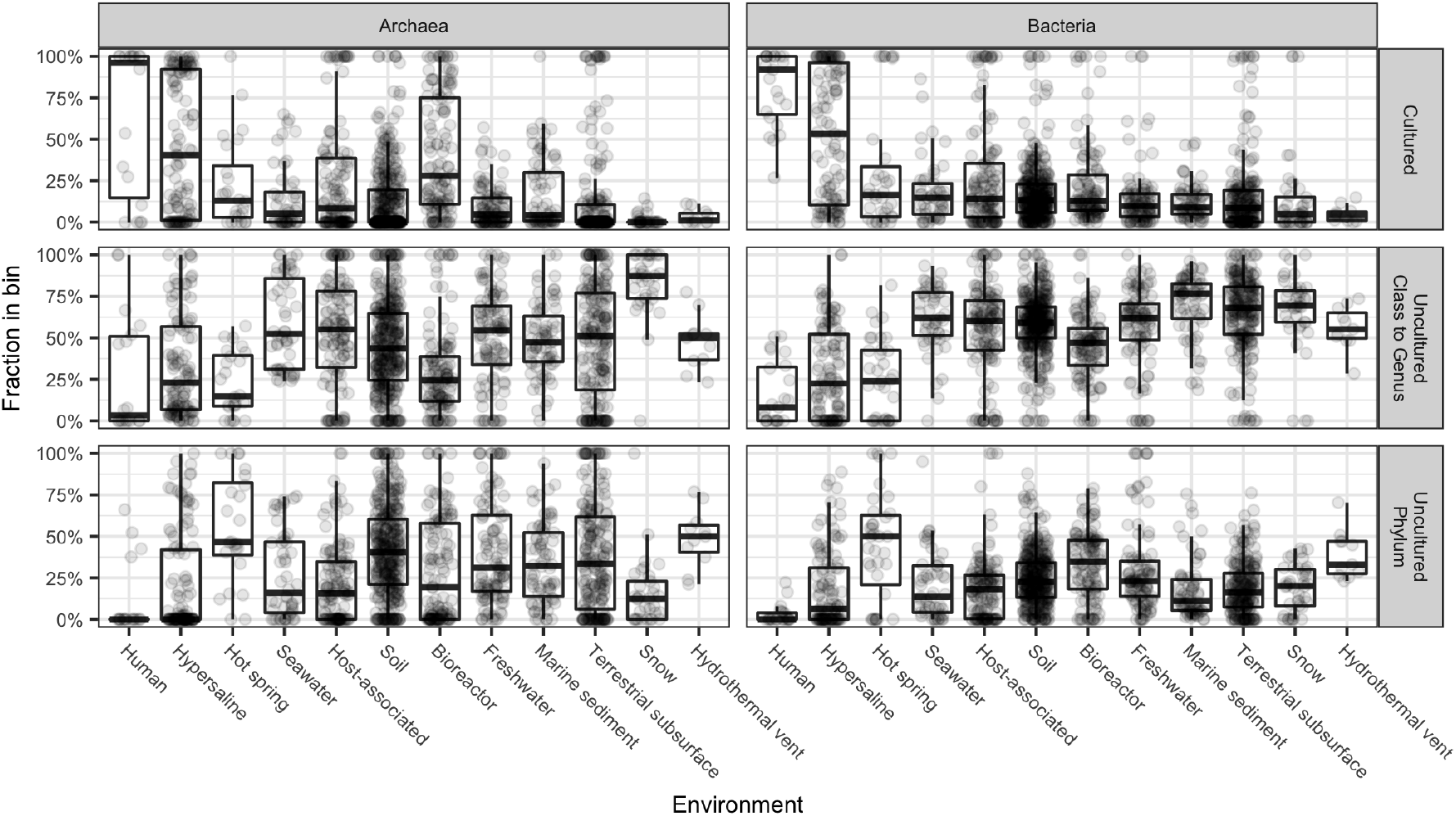
Proportion of 16S rRNA gene sequences by scaffold read depth averaged across all metagenomes. Each single datapoint represents the abundance of reads in that similarity bin from a single metagenome. Rows represent different similarity bins.

By using a large collection of publicly-available sequences that are as complete a sampling as possible, our sequence abundance quantifications can be extrapolated to global cell estimates, although this approach is biased against cells that are less amenable to DNA extraction and under-sampled environments. Copy numbers of 16S rRNA genes per cell can only be determined for completed genomes (means of 3.8 copies/genome for 1657 bacteria, and 1.8 copies/genome for 79 archaeal genomes on IMG https://img.jgi.doe.gov/mer/, March 30, 2018). However, no complete genomes are yet available for uncultured organisms. Applying the 16S rRNA copy numbers for completed genomes to our estimations of total cells would increase our estimates of the abundance of uncultured organisms, since archaea, which we found to be less well-cultured, would be divided by the smaller number. Therefore, we use the conservative simplification of a single 16S rRNA copy number per genome to estimate that 81% of microbial cells on Earth are from uncultured genera or higher (7.3 × 10^29^ cells) and 25% are from uncultured phyla (2.2 × 10^29^ cells) (Table 1). When abundance data are derived from metatranscriptomes, uncultured cells increase to 98% (5.9 × 10^29^), with uncultured phyla contributing 69% (4.2 × 10^29^) (Table 2). If the terrestrial subsurface datasets lack contributions from the ultra-small uncultured cells missed in standard filtering methods (23), or if DNA extraction favors cultured taxa, which may have more easily-lysed cell membranes, then these values are underestimates for the abundance of uncultured cells on Earth.

**Table 1.** Metagenome-based estimates of global microbial cell abundances from uncultured archaea and bacteria, based on 16S rRNA gene sequence read depths. Estimated fractions are in parentheses. Environments with fewer microbial cells were excluded.

| Environment | Cells x 10<sup>26</sup> |  |  |  |
| --- | --- | --- | --- | --- |
|  | Total microbial cells | Cultured species to genera<sup>#</sup> | Uncultured genera to classes<sup>#</sup> | Uncultured Phyla and higher<sup>#</sup> |
| Marine sediment (48) | 2,900 | 390 (13%) | 1,921 (66%) | 590 (20%) |
| Soil (49) | 2,560 | 454 (18%) | 1,268 (50%) | 839 (33%) |
| Terrestrial subsurface (49) | 2,500 | 702 (28%) | 1,211 (48%) | 587 (23%) |
| Seawater (49) | 1,010 | 143 (14%) | 640 (63%) | 229 (23%) |
| Freshwater (49) | 1.3 | 0.1 (11%) | 0.8 (64%) | 0.3 (25%) |
| Plant hosts (50) | 1 | 0.5 (49%) | 0.4 (37%) | 0.1 (14%) |
| Animal hosts (51) | 0.2 | 0.1 (49%) | 0.1 (37%) | 0.0 (14%) |
| **Sum** | **8,974** | **1,689 (19%)** | **5,050 (56%)** | **2,245 (25%)** |

<sup>#</sup>Cut-offs are the upper 95% confidence interval of the median 16S rRNA gene identity for each taxonomic level (13).

**Table 2.** Metatranscriptome-based estimates of global microbial cell abundances from uncultured archaea and bacteria, based on 16S rRNA gene sequence numbers. Estimated fractions are in parentheses. Environments with fewer microbial cells were excluded.

| Environment | Cells x 10<sup>26</sup> |  |  |  |
| --- | --- | --- | --- | --- |
|  | Total microbial cells | Cultured species to genera<sup>#</sup> | Uncultured genera to classes<sup>#</sup> | Uncultured Phyla and higher<sup>#</sup> |
| Marine sediment (48) | NA | NA | NA | NA |
| Soil (49) | 2,560 | 49 (2%) | 758 (30%) | 1,753 (69%) |
| Terrestrial subsurface (49) | 2,500 | 45 (2%) | 597 (24%) | 1,858 (74%) |
| Seawater (49) | 1,010 | 36 (4%) | 389 (38%) | 587 (58%) |
| Freshwater (49) | 1.3 | 0.0 (3%) | 0.5 (40%) | 0.7 (56%) |
| Plant hosts (50) | 1 | 0.2 (18%) | 0.3 (33%) | 0.5 (49%) |
| Animal hosts (51) | 0.2 | 0.0 (18%) | 0.1 (33%) | 0.1 (49%) |
| **Sum** | **6,074** | **129 (2%)** | **1,744 (29%)** | **4,200 (69%)** |

<sup>#</sup>Cut-offs are the upper 95% confidence interval of the median 16S rRNA gene identity for each taxonomic level (13).

We tested whether only a few clades account for this global dominance of uncultured microbes, suggesting that problem of culturability could be solved by just getting a few important species into culture. On the contrary, each category of phylogenetic novelty contained many different genera (Fig. 4). Also, genera at all levels of phylogenetic novelty were distributed throughout the rank abundance curves in all environments except for humans (Fig. 4). The taxonomic identities of the ten most abundant genera differed between environments, and often included genera from newly-named uncultured phyla such as Parcubacteria, Omnitrophica, Latescibacteria, Patescibacteria, Bathyarchaeota, Woesearchaeota, Armatimonadetes, AC1, Miscellaneous Euryarchaeotal Group, Saccharibacteria, WS6, Marinimicrobia, and FBP (Fig.4). Despite having fewer overall sequences than bacteria, archaea were in the ten most abundant genera in 8 of the 12 environments. Few of the top ten genera in metagenomes were also in the top ten genera in metatranscriptomes. The exception was Chloroflexi_Anaerolineaceae, which was present in the top ten genera in both datasets for hot springs, terrestrial subsurface, and bioreactors. However, this could be an artifact, since uncultured members of this group have not been taxonomically characterized to the genus level, so these bins may lump together many different genera collectively labeled as “uncultured”. Some of the most abundant uncultured clades, such as *Candidatus Pelagibacter sp.* in seawater, have actually been obtained in pure culture (24), but their physiological requirements prevent them from meeting stringent requirements, such as the ability to be grown out of stocks of cells preserved in glycerol at −80°C, required to receive an official taxonomy. However, few other examples of such cryptically cultured organisms occur in our dataset.

**Fig. 4.**
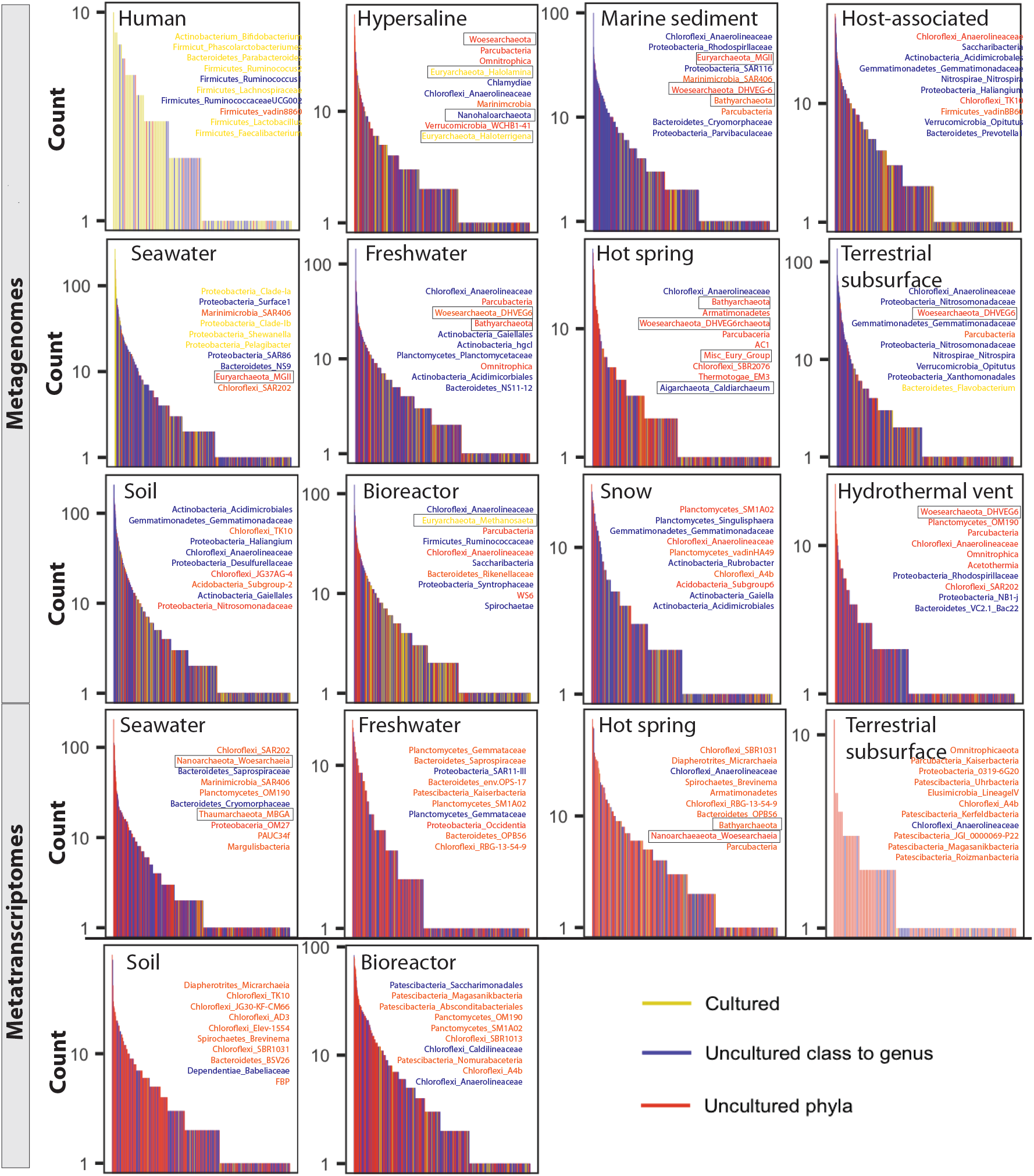
Rank abundance plots by taxonomic genus assignments for metagenomic data (top three rows) and metatranscriptomic data (bottom two rows). Listed in each box are the top ten most abundant genera for that environment in the format of phylum_lowest identified taxonomic group, with archaea outlined in black. Data are colored for uncultured phyla (orange), uncultured class to genus (blue) or cultured (yellow). Taxonomic-based genera that had sequences from multiple phylogeny-based percent identity bins were labeled with the color of the bin with the most sequences.

Many of the top ten genera were taxonomically identified as belonging to cultured phyla, even though we found them to be <86% similar to their nearest cultured neighbor. This is because taxonomic identification and phylogenetic identification are not identical methods. Sequences that have low similarity to cultures can nonetheless be given a taxonomic classification to a cultured phylum because the database used for classification also contains many instances of uncultured sequences that have previously been named as part of that phylum. When genomes become available, such groups are often re-assigned as phyla (1). Our results suggest that rare and abundant taxa are both cultured and uncultured, as well as bacterial and archaeal.

Our datasets likely include some amount of relic preserved DNA that can inflate diversity estimates (25). However, we do not calculate total diversity in a single sample, but rather occurrence frequency across many samples. Extracellular DNA from a particular taxonomic group is not likely to be abundant in the majority of samples, to the exclusion of intracellular DNA from that taxonomic group. In addition, in all environments, metatranscriptomes were characterized by higher fractions of sequences from uncultured groups than the metagenomic databases were, with particularly high contributions from uncultured phyla (Fig. 2). This suggests that the uncultured cells that dominate these datasets likely come from living organisms.

These results offer at least a partial explanation for “the great plate count anomaly”, which states that <1% of environmental microbial cells are culturable with standard methods (26). To update this analysis, we examined 347 experiments in 26 studies from lakes, rivers, drinking water, seawater, marine and terrestrial subsurface, animal hosts, and soils, and found a median of 0.5% culturable cells (Table S3). The past several decades have seen considerable progress on novel culturing techniques, which have yielded higher fractions of culturable cells (25 ± 20%, n = 38) in fish guts (27), rice paddies (28), surface marine sediments (29, 30), agricultural soils (31), and eutrophic lakes (26). However, these studies expanded the set of cultured taxa only to novel families (29, 31), and we show that the percentages of cells from cultured families in these environments match these studies’ percentages of culturable cells (Table S4). Therefore, we propose that these innovative methods likely were successful at culturing viable but non-culturable cells (VBNC), which are cells from previously-cultured cultured clades that are temporarily and reversibly culture-resistant (32). However, our analysis shows that a considerable fraction of cells in non-human environments are phylogenetically divergent, even belonging to novel phyla. We propose that representatives of these phyla resist cultivation due to more fundamental reasons, making them phylogenetically divergent non-cultured cells (PDNC). We roughly define PDNC as cells from orders or higher with no cultured representatives. Unlike VBNC, PDNC are not dormant close relatives of cultured species that can be expected to behave like known cultures if given the correct combination of growth conditions. These entire lineages may have physiologies that prevent growth in pure culture, such as dependences on syntrophic interactions (33), precise chemical or physical parameters that are difficult to maintain (29), extreme dependence on oligogrophy (24, 34, 35), or very slow growth rates (36). Two of the most recent uncultured phyla that have been brought into pure culture are Nitrosopumilus sp., from the Thaumarchaeota phylum (34), and *Abditibacterium utsteinense,* from the FBP phylum (37). These required extremely low nutrient environments, incubation times of many months, and, in the case of *A. utsteinense,* high antibiotic levels to keep out competitors. Fundamentally novel culturing techniques, possibly guided by insights on cell physiology derived from genomic studies, are likely required to get more of these highly abundant and deeply-divergent clades in culture.

Given the substantial functional differences that often exist between closely-related microbial species or strains, these uncultured lineages are likely to contain many novel metabolic pathways, enzyme functions, cellular structures, physiologies (38). For instance, uncultured clades of archaea and bacteria have more genes that are un-annotatable with current databases than cultured clades (27 and 37%, vs. 19 and 31%, respectively, Fig. S1). In addition, a rapidly growing number of studies are uncovering potentially important functions of uncultured clades within specific environmental contexts (20, 39-41).

We conclude that uncultured taxa, often at very high levels of phylogenetic novelty, are abundant and alive in Earth’s microbiome, and may harbor undiscovered functions that are important on an ecosystem level. The high proportion of sequences from uncultured groups in human-maintained bioreactors, animal and plant hosts, and soils, many of which were agricultural or municipal, shows that highly divergent novel clades are not only a feature of pristine wilderness environments, but are important in engineered environments with immediate human applications as well. This suggests that *ex situ* experiments on existing microbial cultures may not represent the functions of the majority of cells *in situ.* For environmentally important VBNC cells, novel culture techniques are showing great success in getting them into culture (35, 42). For PDNC, novel culture-independent techniques such as genomic inference (43), label incorporation (44–46), or tracking slow growth in a mixed population under different conditions (47), will allow the study of their physiology and ecology, and guide efforts to culture them.

## Acknowledgments

This work was supported by (1) the National Science Foundation under grant numbers OCE-1431598 (KL), 1137097 (LC and JY), and 0711134 (LC and JY), (2) the NSF Center for Dark Energy Biosphere Investigations (0CE-0939564) contribution # to be filled in (KL), (3) the Alfred P. Sloan Foundation Fellowship (FG-2015-65399, KL), (4) the Simons Foundation (404586, KL), and (5) NASA Exobiology (NNX16AL59G), (6) the University of Tennessee, Knoxville College of Arts and Sciences (LC and JY), (7) the Joint Institute for Computational Sciences and the Beacon Project (LC and JY), (8) Intel Corporation through an Intel Parallel Computing Center award to support development of HPC-BLAST (LC and JY), and (9) the Oak Ridge National Laboratory Science Alliance. Andrew D. Steen helped with the R code and Tatiana Vishnivetskaya performed the Russian to English translations. Any opinions, findings, conclusions, or recommendations expressed in this material are those of the author(s) and do not necessarily reflect the views of the National Science Foundation, the University of Tennessee, or Intel Corporation.

## Author Contributions

K. G. L. conceived of and obtained funding for the project, analyzed data, and led the writing of the paper; J. L. and A. D. S. provided supplementary data analysis and helped write the paper; J. Y. parallelized the BLAST analyses and helped write the paper; L. C. obtained funding, advised J. Y., and helped write the paper.

